# *Nebulosa* recovers single cell gene expression signals by kernel density estimation

**DOI:** 10.1101/2020.09.29.315879

**Authors:** Jose Alquicira-Hernandez, Joseph E. Powell

## Abstract

**Summary:** Data sparsity in single-cell experiments prevents an accurate assessment of gene expression when visualised in a low-dimensional space. Here, we introduce *Nebulosa*, an R package that uses weighted kernel density estimation to recover signals lost through drop-out or low expression.

**Availability and implementation:** *Nebulosa* can be easily installed from www.github.com/powellgenomicslab/Nebulosa

## Main text

Current single cell sequencing technologies allow transcriptional profiling from thousands to millions of individual cells in a single experiment. However, single-cell gene expression data commonly exhibits dropout events derived from stochastic transcription (Ochiai *et al*., 2020), low abundance of transcripts (Kharchenko *et al*., 2014), or shallow sequence depth (Haque et al., 2017). All of these factors affect the visualisation of gene expression in low-dimensional representations such as UMAP, PCA, or *t*-SNE. Such visualisation is crucial in any single-cell data analysis, and is a commonly used approach for cell annotation (based on canonical markers), the discovery of new cell sub-types, and the evaluation of confounding or batch effects.

Here, we introduce *Nebulosa*, a novel visualisation approach to resolve sparsity based on gene-weighted kernel density estimation. We show that by incorporating the cell density information from a low-dimensional space, the gene expression signal from dropped-out genes can be rescued. We applied Nebulosa in two datasets: 68k peripheral blood mononuclear cells (PBMCs) (Zheng *et al*., 2017) and keratinocytes from a mouse transgenic for the E7/E6 genes of the Human Papilloma Virus 16 (HPV16) (Lukowski *et al*., 2018). We first highlight the performance of *Nebulosa* using CD4 expression in PBMCs - a gene that exhibits a high dropout rate, and is predominately restricted to non-cytotoxic T cells and myeloid cells. Visualizing the expression of CD4 in cells using a traditional UMAP, shows an absence of expression, which can be attributed to the large number of cells with dropped-out expression combined with over-plotting (Figure 1a). When plotting cells ordered by gene expression, some of the cells expressing CD4 become more observable (Figure 1b). However, this expression lacks a clear specificity across the UMAP space, resulting in a noisy signal difficult to associate to a given sub-population of cells. Applying Nebulosa to CD4 expression, we observe the clear expression signal in two well-defined clusters corresponding to non-cytotoxic T cells and myeloid cells (Figure 1c). This is achieved by the kernel function smoothing cell density weighted by the gene expression. Nebulosa also removes the localised expression of CD4 from areas where this gene is expressed in very limited cell numbers. These instances are caused by stochastic transcription of other genes, or ambient RNA present in droplets, making biological interpretation difficult. This feature produces a noise-free representation of the gene expression while recovering the signal from cells that are more likely to express a gene based on their neighbouring cells.

**Fig. 1.**
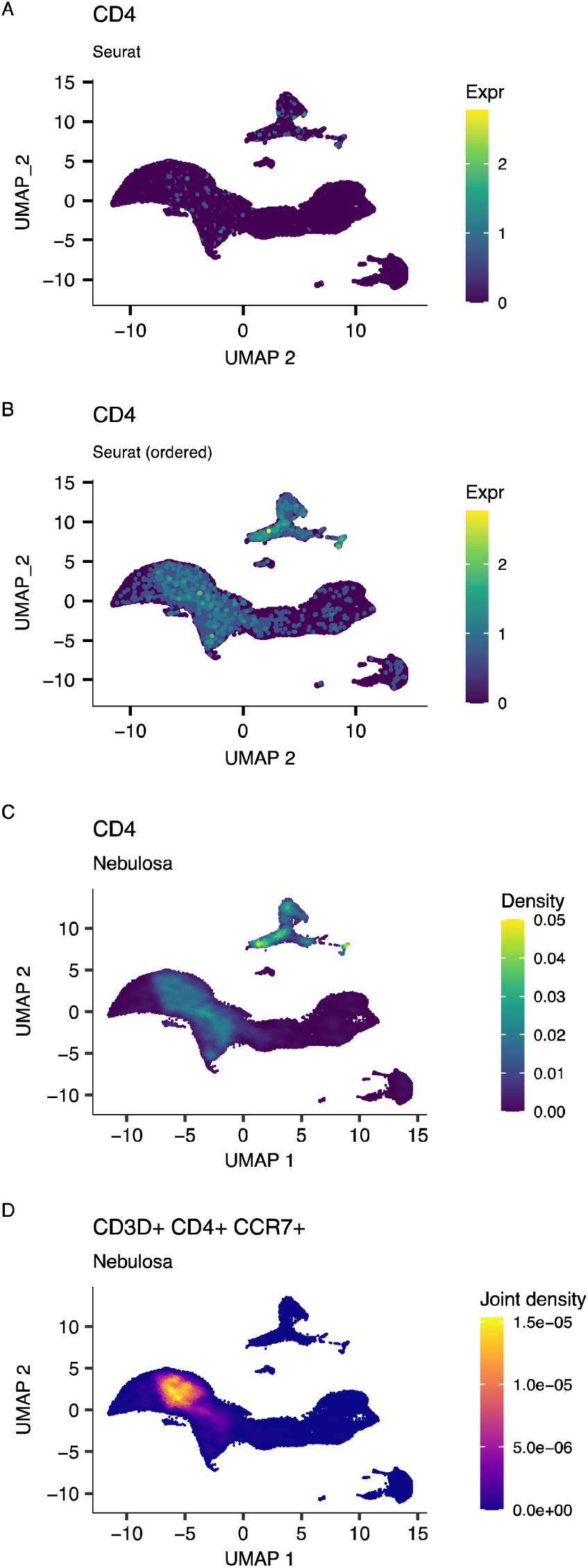
Nebulosa recovers cell gene expression signals that are lost through drop-out or low expression. Cells expressing one or more transcripts of CD4 are plotted for each of 68,000 peripheral blood mononucleated cells using the standard UMAP plotting (A); with cells ordered based on their expression of CD4 (B); and with Nebulosa’s kernel function utilising low-dimension (UMAP) cellular density features. We also highlight the ability to identify cell populations based on joint density estimation of multiple gene markers, using CD3D, CD4, and CCR7 to identify CD4+ cells in peripheral blood mononucleated cells (D)

*Nebulosa* can also create a joint density estimate to visualise the expression overlap of multiple genes. As a demonstration, we used *Nebulosa* to identify naive CD4 T cells based on the joint expression of CD3D, CD4, and CCR7 (Figure 1d). This feature allows the direct and more precise identification of cell types based on the combined expression of various markers. To further demonstrate the utility of this function, we applied *Nebulosa* to detect the expression of the viral oncogenes E6 and E7 from HPV-transgenic keratynocytes. The combined expression of E6 and E7 was observable in two clusters characterised by the expression of Krt5 (see Supplementary Figure 1), that are difficult to identify with traditional UMAP plotting.

*Nebulosa* makes use of weighted kernel density estimation methods (Duong, 2007; Venables and Ripley, 2002) to represent the expression of gene features from cell neighbours. Besides counts and normalised gene expression, it is possible to visualise metadata variables (such as batch, condition, time point etc.), and feature information from other assays (such as CITE-seq protein markers, ATAC-seq signals). *Nebulosa* can take Seurat or SingleCellExperiment objects as input, enabling easy integration into current single cell analysis workflows. The weighted kernel density estimation is calculated as following:

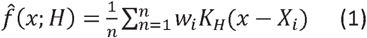

where *n* is the total number of cells, *w*_*i*_ is the scaled gene expression and *X*_*i*_ the embeddings of the *i*_*th*_ cell, *K*(*x*) is a gaussian kernel function. *H* corresponds to the bandwidth matrix for smoothing determined with an unconstrained plug-in selector implemented in the *ks* package and *x* is the density estimate at a given point of the embedding space defined by the grid size used for the computation.

Overall, *Nebulosa* is a powerful approach for visualising single-cell data as it resolves data-sparsity by using the information from neighbouring cells while successfully dealing with over-plotting. *Nebulosa* is easy to use and can be implemented into current standard single cell workflows such as Seurat and Bioconductor workflows.

## Supplementary information

Supplementary data are available online.

## Acknowledgements

We thank Drew Neavin for her suggestions and discussions on data visualization

