## supporting figure 1 for "*Nebulosa* recovers single cell gene expression signals by kernel density estimation"

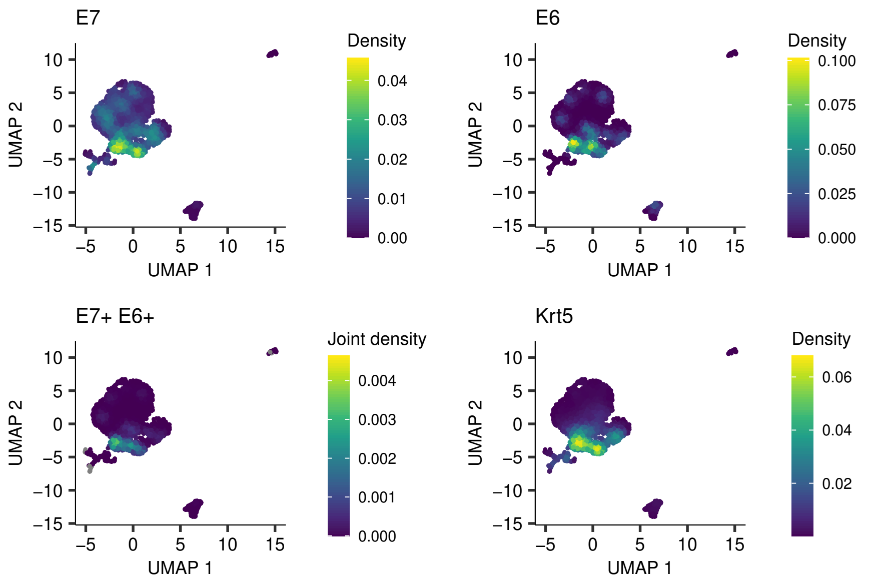


Supplementary Figure 1: Visualization of E7 and E6 gene expression from transgenic mice. Krt5 colocalizes with the E7+ E6+ joint density in keratinocytes.
